# Multi-Scale Coarse Grained Model for the Stepping of Molecular Motors with application to Kinesin

**DOI:** 10.1101/2021.04.05.438476

**Authors:** Yonathan Goldtzvik, D. Thirumalai

## Abstract

Conventional kinesin, a motor protein that transports cargo within cells, walks by taking multiple steps towards the plus end of the microtubule (MT). While significant progress has been made in understanding the details of the walking mechanism of kinesin there are many unresolved issues. From a computational perspective, a central challenge is the large size of the system, which limits the scope of time scales accessible in standard computer simulations. Here, we create a general multi-scale coarse-grained model for motors that enables us to simulate the stepping process of motors on polar tracks (actin and MT) with focus on kinesin. Our approach greatly shortens the computation times without a significant loss in detail, thus allowing us to better describe the molecular basis of the stepping kinetics. The small number of parameters, which are determined by fitting to experimental data, allows us to develop an accurate method that may be adopted to simulate stepping in other molecular motors. The model enables us to simulate a large number of steps, which was not possible previously. We show in agreement with experiments that due to the docking of the neck linker (NL) of kinesin, sometimes deemed as the power stroke, the space explored diffusively by the tethered head is severely restricted allowing the step to be in a tens of microseconds. We predict that increasing the interaction strength between the NL and the motor head, achievable by mutations in the NL, decreases the stepping time but reaches a saturation value. Furthermore, the full 3-dimensional dynamics of the cargo are fully resolved in our model, contributing to the predictive power and allowing us to study the important aspects of cargo-motor interactions.

## Introduction

Molecular motors, or motor proteins, are an integral part of the machinery of living cells. Their most common function is the transportation of cargoes along actin and microtubule (MT) filaments within the cell.^1–3^ To do so, molecular motors convert chemical energy, stored in the form of ATP, into mechanical work. How this is accomplished in a variety of of structurally unrelated motors is the focus of a large body research.^4^

Due to several remarkable single molecule studies of motor proteins performed over the last three decades, we know a great deal about how molecular motors walk along the polar tracks.^5–10^ Furthermore, an increasing number of structures for these motors in various nucleotide states have provided great insights into the inner workings at the molecular level.^11–16^ Nevertheless, understanding the molecular details of how these machines “walk” along cellular filaments continues to pose significant challenges. In particular, due to the spatial and time resolution limitations of current experimental methods, it is difficult to decipher the specifics of the walking mechanism at a microscopic resolution, which could be overcome by using reliable computational models.

The use of Coarse Grained (CG) models in computer simulations has been remarkably successful in elucidating the stepping mechanisms of a variety of motors and other molecular machines.^17–23^ Such models have proven to be particularly useful in dissecting a number of aspects of motor protein motility.^24–27^ The advantage of CG simulations is most evident when considering the relatively large size of molecular motors. The motor domain of kinesin, the smallest of the filament associated motors, is over 300 amino acids long.^11^ Therefore, the simulation of the full motor construct for time scales relevant to the function of the motor are inaccessible to fully atomistic computational methods, necessitating the use of simulations using well-calibrated CG models.

As we alluded to earlier, molecular motors couple the hydrolysis of ATP molecules to the mechanical step taken by the motor.^28,29^ The ATP chemical cycle is a process that takes place at the level of individual protein domains within the motor, involving subtle allosteric transitions.^13,30^ This type of process lends itself to simulations using CG models at the single amino acid resolution.^25,27,31^ The walking or stepping mechanism itself, on the other hand, involves the motion of large and relatively rigid domains.^5,6,8,9,26^ It is, therefore, prudent to construct CG models at the scale of individual protein domains so that multiple steps could be simulated to complement experiments.

In this study, we focus on kinesin, which is perhaps the best known and most researched of all the motor proteins. Kinesin consists of two identical motor domains, connected to each other through flexible coils called the neck linkers (NL).^11,12^ The NLs join to form a long coiled coil structure, named the stalk, which in turn, connects the dimeric motor to the cargo.

Experiments have shown that kinesin walks on MT by a hand-over-hand mechanism. ^6^ According to this mechanism, the motor domain that lies closer to the minus end of the MT, the trailing head (TH), detaches from the MT. Following the TH detachment, the NL of the bound motor domain, the leading head (LH), docks to the LH and propels the detached TH forward toward the plus end. The docking of the NL to the LH, proposed by Vale and coworkers, is some times referred to as the power stroke^32^ in kinesin. The step ends with the TH binding to its target binding site (TBS) on the MT, making it the new LH, and the cycle is repeated. This results in kinesin taking multiple steps (referred to as processive motion) before disengaging from the MT.

Our previous studies of the kinesin step using residue level CG models have been successful at revealing some of the molecular details of a single step.^26,33,34^ Nevertheless, due to the large size of the system, such models are limited in their scope, and extending them to simulate multiple steps is difficult from a computational point of view. To circumvent this problem, we have developed a new multi scale CG (MSCG) model of the kinesin motor. The model consists of a combination of rigid body representations of certain parts of the motor and flexible multi particle representation of the rest of the system. Using this approach the simulation speed is enhanced by approximately 100 fold, which allows us to better explore the parameter space of our model as well as the intrinsic mechanical aspects of the kinesin motor itself.

## Methodology

### Coarse Grained Model

We use multiple levels of coarsening to create a model for simulating the stepping mechanism of the kinesin motor. We model the motor domain using 39 beads of varying radii (See Fig.1). Each bead represents a group of amino acids that are in the same region (spatially close) of the protein structure. The choice of amino acid groups is somewhat arbitrary, with the guiding principle being that the overall shape of the motor domain should be maintained while using a relatively small number of beads (the details of the amino acid groupings is given in the Supplementary Information (SI)). In order to construct the MSCG representation, we use the known crystal structure of the motor domain (Protein Data Bank (PDB) ID: 2KIN). The position of each bead is given by the geometrical mean of the *C*_*α*_ carbon positions of the amino acids represented by that bead. The bead radius is taken to be the radius of gyration of the group of associated *C*_*α*_ carbon atoms.

**Figure 1:**
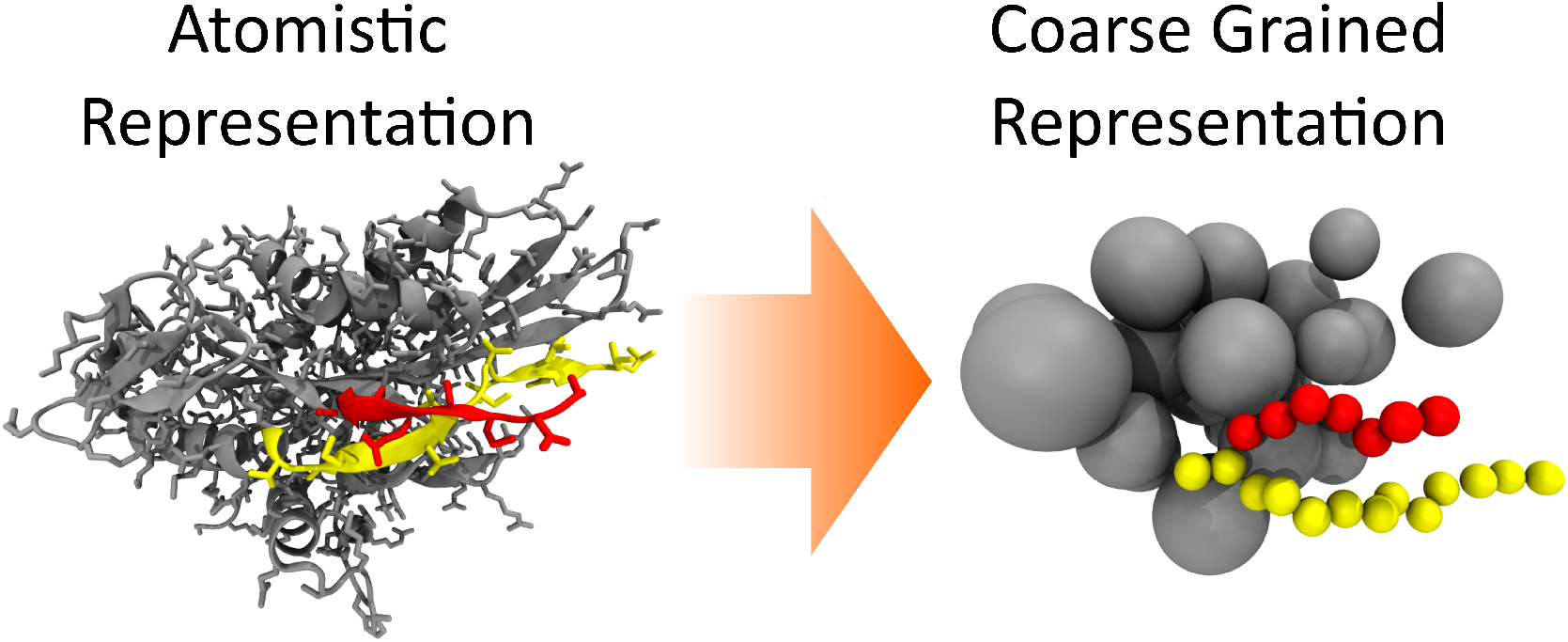
A visual representation of the coarse graining of the kinesin motor domain. On the left is the fully atomistic representation of the protein, and on the right is the MSCG representation, with a much smaller number of beads.

The kinesin motor domain has two flexible structural elements: the Cover-Neck Bundle (CNB) at the N-terminus (red in Fig.1), and the NL at the C-terminus of the motor domain (yellow in Fig.1). Due to their relatively flexible nature, we represent each amino acid in these two structural elements using a single bead, centered at the *C*_*α*_ carbon, with a radius of *σ*_*AA*_ = 1.9Å.

The MSCG representation of the stalk consists of a series of linked beads with a radius *σ*_*S*_ = 5.0Å, which is approximately the radius of the stalk. We use a 30 bead long stalk in the simulations. However, this number could be changed arbitrarily to model stalks of different lengths. We represent the cargo as a single large bead that is attached to the end of the stalk. The radius of the cargo is taken to be *σ*_*MT*_ = 0.5*µm*, which is the typical size used in experiments. ^35–37^ Finally, we represent the MT as a cylinder with a radius of *σ*_*MT*_ = 12.7*nm* (127Å), with its main axis of symmetry lying along the x axis. The complete structure of the kinesin dimer and stalk (excluding the cargo) in the MSCG representation is shown in Fig.2.

**Figure 2:**
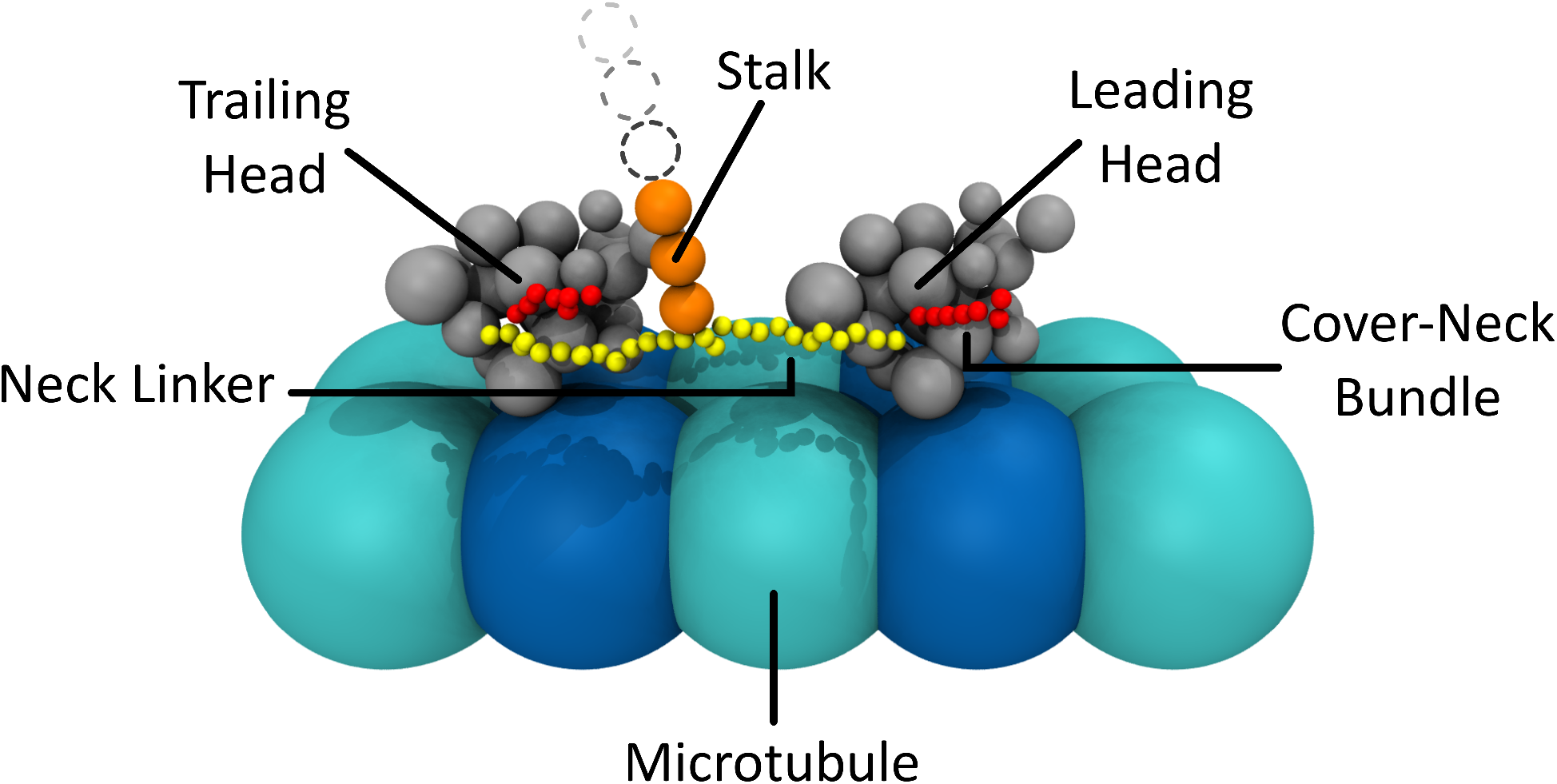
The MSCG representation of the full system (excluding the cargo). The two motor domains (gray spheres) are the TH on the left and LH on the right. The CNB and NLs are represented in red and yellow, respectively. The stalk is depicted by the orange beads. The first orange bead, attached to the NL, is referred to as the Neck Linker Stalk Junction (NLSJ). Finally, the MT is represented by the blue and cyan spheres.

#### Rigid Body Formulation

The basis for our model lies in the observation that the kinesin motor domains maintain a relatively rigid shape during the stepping process, and do not undergo major conformational changes. This allows us to approximate them as rigid bodies. For each rigid element, we assign an Object Frame of Reference (OFR). The position of the rigid body (which corresponds to the OFR origin) with respect to the Laboratory Frame of Reference (LFR) is given by a 3-dimensional Cartesian vector. The rigid body OFR also specifies the orientation of the object, which can also be represented by a 3-dimensional vector. Therefore, the rigid body can be fully described by a 6-dimensional vector,

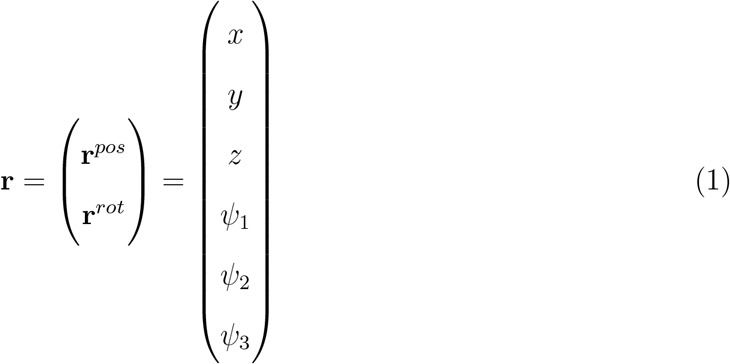

where *x, y*, and *z* are the components of **r**^*pos*^, the position vector, and *ψ*_1*−*3_ are the components of **r**^*rot*^, the rotation vector. We use **r** to mean **r**^*pos*^ by default unless we indicate otherwise.

Once the OFR is assigned, one can construct the components of the rigid body within the OFR. Our model contains four different rigid elements: two motor domains (TH and LH), the NL and Stalk Junction (NLSJ), and the cargo. For the motor domains, the positions of the 39 beads with respect to the OFR are given by 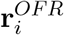 (*i* ∈ {1...39}). The bead positions with respect to the LFR are given by,

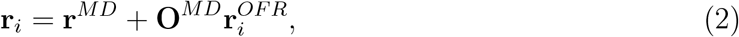

where **r**^*MD*^ is the position of the motor domain rigid body with respect to the LFR, and **O**^*MD*^ is a 3 × 3 transformation matrix that corresponds to the orientation of the OFR with respect to the LFR and is a function of **r**^*rot*^ (if **r**^*rot*^ is a Cartesian rotation vector, **O**^*MD*^ can be obtained using Rodrigues’ rotation formula, which is described clearly by Valdenebro^38^). Both the NLSJ, which is the first bead of the stalk (Fig. 2), and the cargo, are each represented by a single bead with its OFR being at the center. The rest of the system components (CNB, NL, and stalk) are described by simple isotropic beads, and do not require the definition of an OFR.

#### Energy Functions

A major advantage of any rigid body formulation is that the internal interactions within the rigid body are included implicitly within the rigid constraints, thus greatly reducing the computation times. It is, therefore, sufficient to describe their external interactions with the other components of the system. We define the total energy function for the system, *U*_*T*_, as,

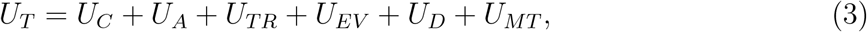

where the different subscripts denote the contributions from connectivity (C), angular (A), torsion (TR), excluded volume (EV), docking (D), and MT interactions.

#### Connectivity Potential

The connectivity term *U*_*C*_ accounts for non-breakable chemical bonds in the system. The functional form of the potential is given by the Finite Elastic Non Extensible (FENE) potential,

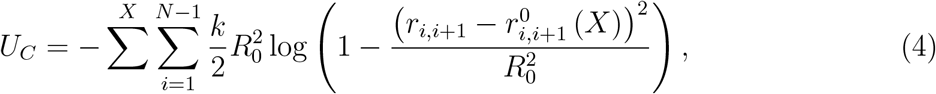

where 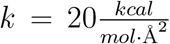 and *R*_0_ = 2Å. The equilibrium distance, 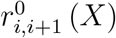, depends on the component of the motor,

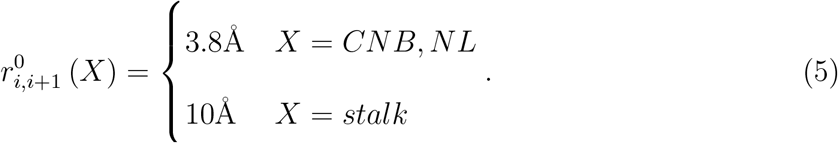

To account for connectivity between the CNB and NL to the motor domain, we treat the last (first for NL) bead in each of these chains as if it is a part of the rigid element. In other words, the bead position obeys Eq.2 and its position within the OFR is obtained from the PDB structure. Similarly, the last beads of the two NLs are connected to the NLSJ by the same constraint as is the case for the last bead in the stalk and the cargo. The positions of the last beads of the NLs within the OFR of the NLSJ are at **r** = (±2, 0, −6.6) in Å. The position of the last bead of the stalk within the OFR of the cargo is at the cargo surface with **r** = (0, 0, −0.5) in *µm* (see Fig.4).

**Figure 3:**
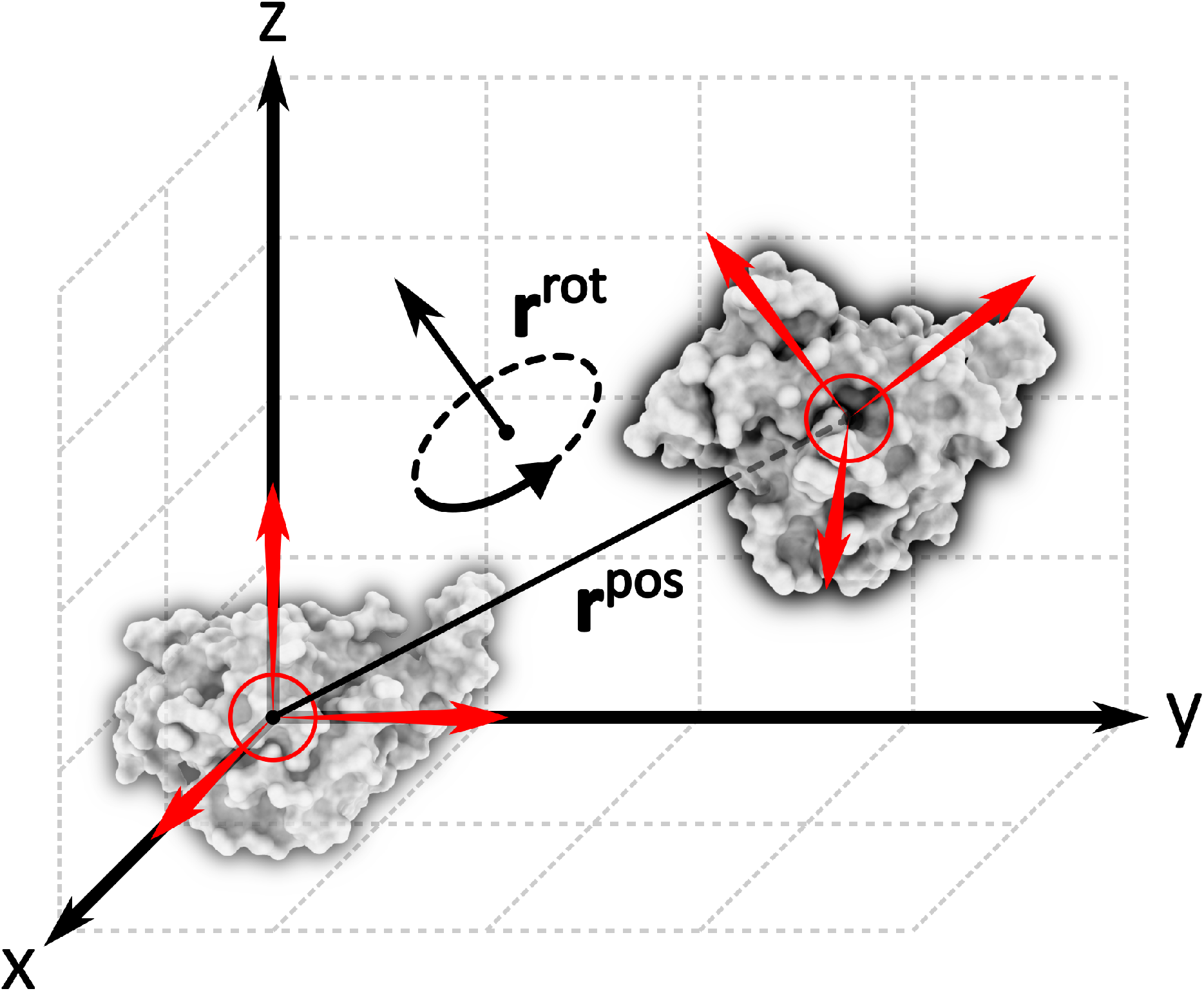
Schematic of the transformations of the OFR (Object Frame of Reference) relative to the LFR (Laboratory Frame of Reference). The position of the OFR origin is given by **r**^*pos*^ while the orientation of the OFR relative to the LFR is given by **r**^*rot*^.

**Figure 4:**
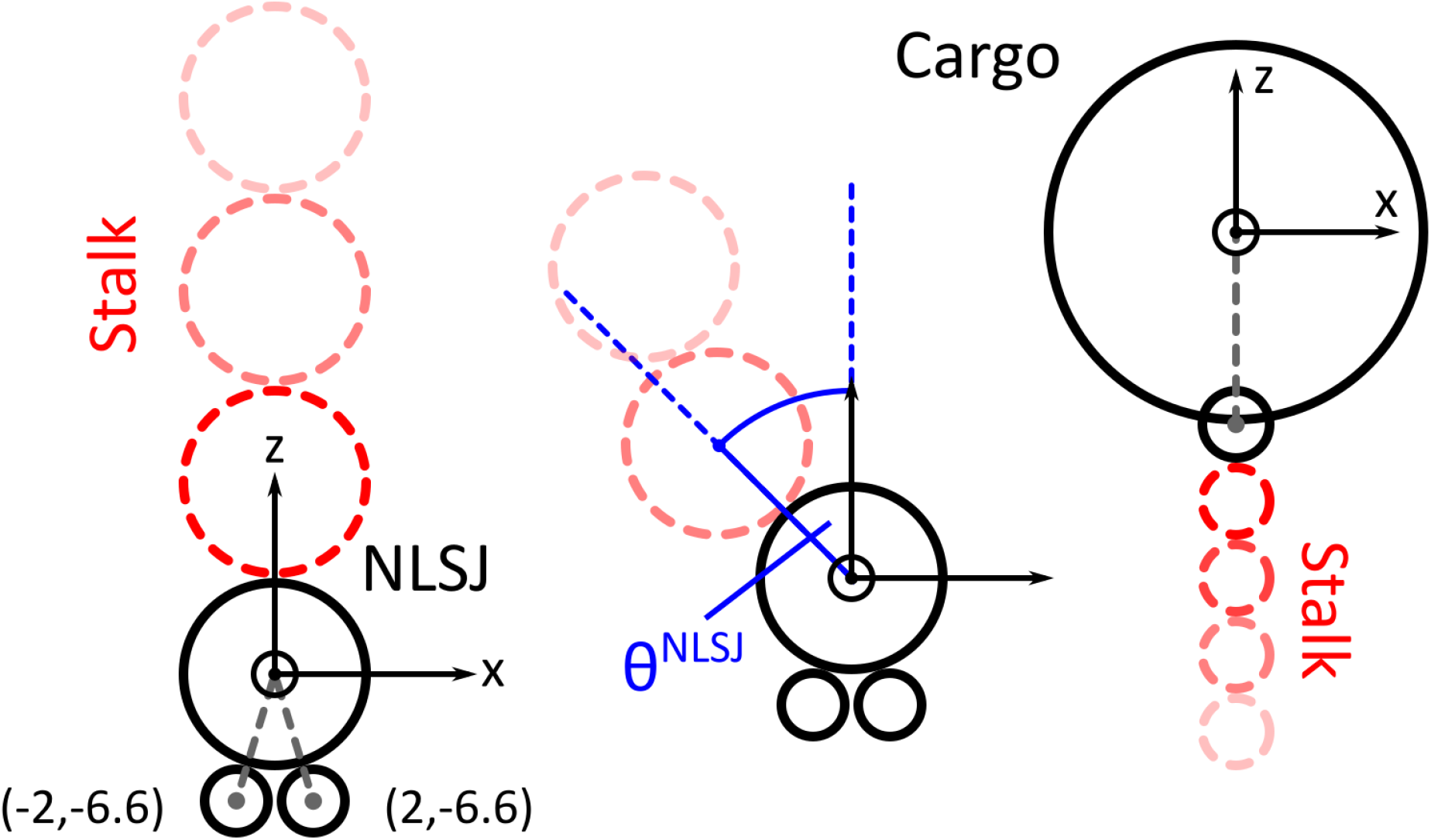
Left: The positions of the contact points of the NLs with the NLSJ within the OFR of the NLSJ. Middle: The definition of *θ*_*NLSJ*_ in terms of the orientation of the stalk with respect to the axes of the OFR of the NLSJ. Right: The position of the contact point between the stalk and the cargo within the cargo OFR.

#### Angular Potential

To ensure that the stiffness of the stalk is properly maintained, we apply an angular potential,

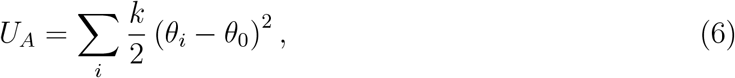

where 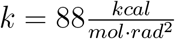 and *θ*_0_ = *π*, and the angle *θ*_*i*_ is the angle formed by any three consecutive beads *i, i*+1, *i*+2. The summation is over all such angles along the stalk, including the NLSJ and the cargo. The value of the stiffness parameter *k* was chosen such that the persistence length of the stalk is approximately 150nm which is a typical value for coiled coil structures. ^39^

Since the ORF of the NLSJ has to be properly aligned with the rest of the stalk, we added an additional term of the form 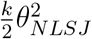 to *U*_*A*_. In this case, *θ*_*NLSJ*_ is the angle between the z axis of the NLSJ OFR and the line connecting the center of the NLSJ and the next bead in the stalk (see Fig.4).

#### Torsional Potential

Interactions between the cargo and the motor are mediated by the stalk. More specifically, when both the motor domains are bound to the MT, the motor imposes constraints on the motion of the cargo. The most obvious constraint is on the translation of the cargo, which forces the cargo to stay within a certain distance from the motor. In our model, this constraint manifests itself through the connectivity term (and to some extent the angular term) of the energy function. An additional, and less obvious constraint is the one on the rotation of the cargo. It is well established that, when both the motor domains are bound to the MT, the motor imposes a rotational constraint on the cargo through torsional strain along the stalk. ^40,41^ As a result, the cargo is not entirely free to rotate along its z axis.

To account for this effect, we introduce a torsional term, *U*_*TR*_, in the energy function, that couples the orientations of the NLSJ and cargo OFRs:

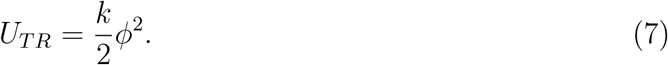

The angle *ϕ* reflects the torsional shift between the base of the stalk (NLSJ end) and the top of the stalk (cargo end). Since we represent the stalk using isotropic beads, the torsional information is lost. To approximate the torsional shift, we make the following calculation. The torsional shift should be reflected in the relative rotations of the NLSJ and the cargo along the z axes of their respective OFRs. In other words, the torsional shift should be equal to the angle between the x axes (or any unit vector within the xy plane of the OFR) of the two rigid objects. The problem is that the xy planes of the two OFRs are not necessarily parallel to each other at any given instant, as both the NLSJ and the cargo rotate 3-dimensionally in space. To circumvent this problem, we perform a virtual rotation of the NLSJ OFR that aligns its z axis with the cargo z axis (see Fig.5). The axis for this rotation transformation is simply 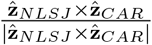. Once the two z axes are aligned, we can obtain the angular difference between the two x axes easily. It is important to note that the domain of *ϕ* is not limited to (−*π, π*), since the cargo can in theory perform multiple rotations. The spring constant *k* takes different values, depending on the desired torsional strain.

**Figure 5:**
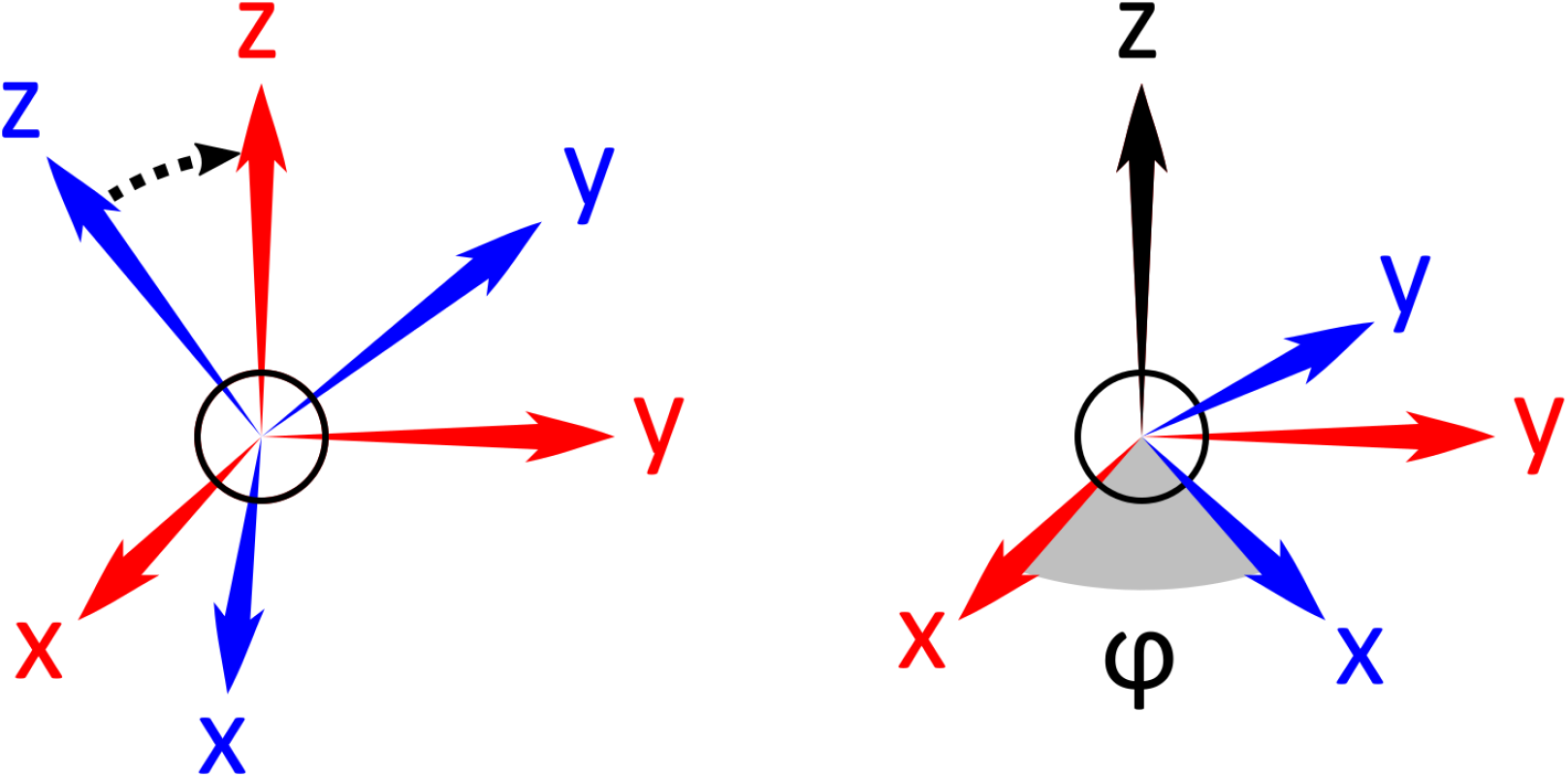
The torsional angle *ϕ* is obtained by first aligning the z axes of the OFRs of the cargo and the NLSJ (left). Once the axes are aligned, *ϕ* is given by the angle between the corresponding axes in the two OFRs (right).

#### Excluded Volume Potential

To account for the repulsive interactions between the different beads in our model, we use the following expression for the excluded,

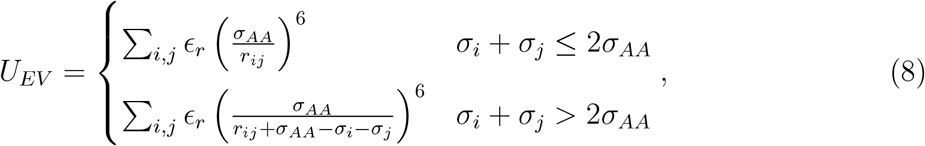

where 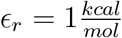 is the distance between beads *i* and *j, σ*_*i*_ and *σ*_*j*_ are the radii of beads *i* and *j* respectively, and *σ*_*AA*_ = 1.9Å is the typical radius of an amino acid. The summation is over all pairs of particles in the system. The reason for choosing the form of excluded volume potential given in Eq.8, is to ensure a uniform range of interactions for beads of varying sizes.

Excluded volume interactions involving the MT contribute an additional term to *U*_*EV*_,

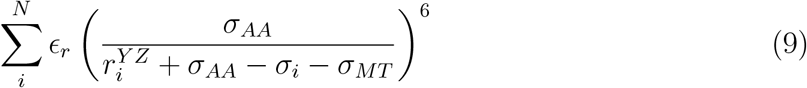

where 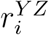 is the length of the projection of the bead position vector **r**_*i*_ onto the YZ plane. The summation is over all the particles in the system.

#### Docking Interactions

As we discussed earlier, the NL of the LH undergoes a docking transition upon ATP binding to the LH, which is deemed the power stroke. This conformational change is fundamental to the stepping mechanism, as it propels the detached TH froward toward the TBS, and restrains the motor to step predominantly on a single protofilament on the MT.^34^ When undocked, the NL is unfolded and we therefore treat it simply as a self-avoiding chain. In order to induce docking, we introduce attractive interactions between the NL and the LH motor domain, given by,

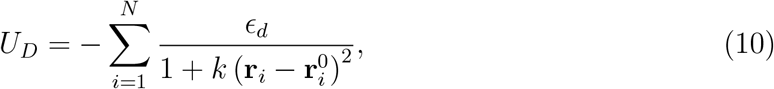

where 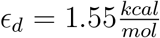 is the interaction energy, and *k* = 0.07Å^2^ determines the stiffness of the potential. The choice of *k* is made so that when 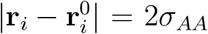, the energy is ≈ 0.5 *ϵ*_*d*_ in Eq.10. **r**_*i*_ is the position of the NL bead, and 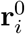 is the position of the center of the energy well. 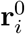 is determined by the position of the docked bead in the crystal structure (PDB ID: 2KIN) of the kinesin motor domain with a docked NL. It may be thought of as the position of a pseudo-bead in the motor domain rigid body, and therefore, it obeys Eq.2 with 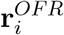 as its position in the OFR. These interactions were applied both to the NL and the CB in the docked state. The choice of *ϵ*_*d*_ will be discussed below.

#### MT Interactions

Generally speaking, kinesin can be found in one of two states in terms of affinities for the MT. In the *apo* and ATP bound states, kinesin binds MT with high affinity, while when bound to ADP, kinesin’s affinity for MT is significantly decreased. The interactions between the motor domain of kinesin and the TBSs on the MT are described by *U*_*MT*_ (Eq.3), which has a similar form to Eq.10. The energy parameter *ϵ*_*d*_ is replaced with *ϵ*_*MT*_, which takes one of two values, depending on the chemical state of the motor domain: 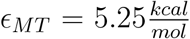 when the motor domain is in the strongly bound state, and 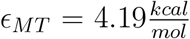 when the motor domain is in the weakly bound state (see below for these choices). The sum is over the 6 beads that are at the interface between the MT and the motor domain in the CG representation (see Fig.S2 and the SI for details). 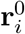 in this case represents the positions of the beads when bound to the MT. To obtain these positions and the correct orientation of the motor with respect to the MT, we aligned the crystal structure of kinesin (PDBID: 2KIN), which were used to construct the CG model, with the structure of a kinesin motor domain bound to MT (PDBID: 2P4N).

### Simulation Details

#### Brownian Dynamics

We assume that the dynamics of the kinesin step are governed by Langevin dynamics in the over-damped limit. Therefore, we can simulate the motion of the system using the Ermak-McCammon Brownian dynamics algorithm, ^42^

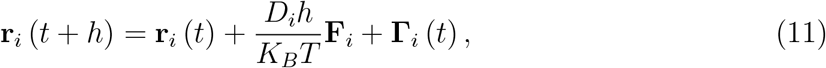

where **r**_*i*_ (*t*) is the position of bead *i* at time t, *h* is the time step, *D*_*i*_ is the diffusion coefficient of the bead, **F**_*i*_ is the systematic force acting on the bead, and **Γ**_*i*_ (*t*) is a random force acting on the bead at time t.

We assume that the diffusion coefficient obeys the Stokes-Einstein equation, 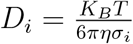, where *σ*_*i*_ is the bead radius, and *η* is the viscosity. As required by the fluctuation-dissipation theorem, the random force obeys ⟨**Γ**_*i*_ (*t*) ⟩ = 0 and ⟨**Γ**_*i*_ (*t*) · **Γ**_*i*_ (*t′*) ⟩= 6*D*_*i*_*hδ*_*tt′*_. In the implementation of the algorithm, we used the BAOAB method that has been shown to have better convergence when compared with the standard Euler-Maruyama method.^43^

#### Rigid Body Brownian Dynamics

To address the dynamics of the rigid elements we used an adaptation of Brownian dynamics.^44^ We first define the generalized coordinates of the ORF orientation. The orientation of a rigid object can be obtained by rotating a reference object around a certain axis. We therefore define **r**_*rot*_ as a Cartesian rotation vector that lies along the rotation axis whose size is equal to the rotation angle in radians. Cartesian rotation vectors are extremely useful in describing 3-dimensional rotations, and as we stated earlier, it is easy to find their corresponding transformation matrices, *O*, using Rodrigues’ rotation formula.

A disadvantage of the Cartesian rotation vectors as a generalized coordinate is that it is difficult to obtain an expression for a rotation as a function of two previous rotations. To circumvent this problem we use the Gibbs rotation vector, **g**_*rot*_.^45^ This vector is similar to the Cartesian rotation vector, but, given that *θ* is the rotation angle, its length is equal to tan 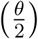 instead of *θ*. The advantage of the Gibbs representation of rotation is that there is a simple formula for the summation of two consecutive rotations, 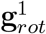 and 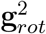, given by,

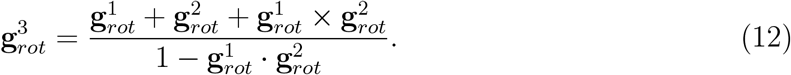

Since we are interested in using the Cartesian representation, we introduce a conversion operator, 𝒢 (**r**_*rot*_) = **g**_*rot*_, and its inverse, 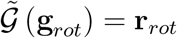, and define the following operator,

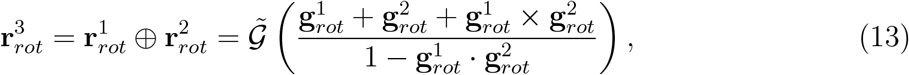

where 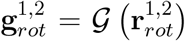. In other words, the operator ⊕ is the rotational equivalent of addition in terms of Cartesian rotation vectors. This approach is based on the one introduced by Antosiewicz *et al* .^44^ An alternative approach that is commonly employed is the use of quaternions introduced for use in molecular simulations of polyatomic molecules.^46^ Both approaches are equally valid and do not affect the results of the study.

Having introduced the generalized coordinates, we can define our algorithm for the rigid body Brownian dynamics. Since we are now dealing with an anisotropic rigid body, the diffusion coefficient, *D*, has to be replaced with a diffusion tensor, **D**. Because the translation and rotation of rigid bodies are not necessarily independent, **D** is a 6-dimensional square matrix that couples both the force and torque acting on the object. In analogy to the 6-dimensional vector **r** in Eq.1, we define a 6-dimensional generalized force,

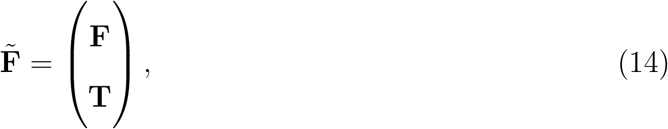

where **F** and **T** are the components of the force and torque, respectively. The force component is calculated by summing over all the individual forces acting on the different beads in the rigid body, 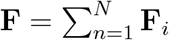. Similarly, the associated torque is calculated by summing over all the torques that result from these individual forces, 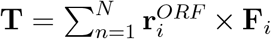. To obtain the coupled generalized force, we simply multiply 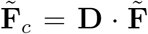. We refer to the resulting force and torque as **F**_*c*_ and **T**_*c*_ respectively.

Similarly, we define a 6-dimensional random noise vector,

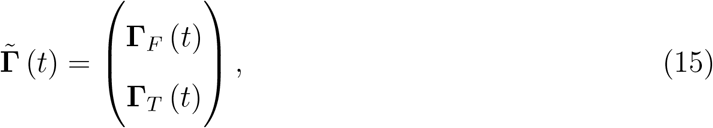

with a force component **Γ**_*F*_ (*t*), and a torque component **Γ**_*F*_ (*t*). Just like the 3-dimensional noise, the random generalized force vector obeys 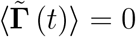 and 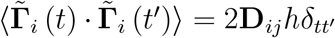.

Once the forces and torques are coupled, we use the following algorithm to update the positions and rotations,

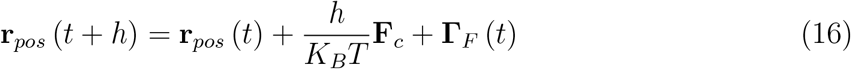

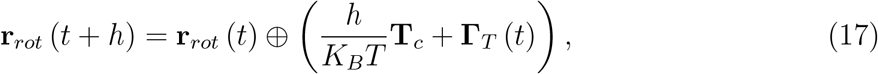

where the ⊕ symbol is defined in Eq.13. This integration scheme is applied to every rigid object in the system, which are, the two motor domain, the NLSJ, and the cargo.

#### Diffusion Tensor Calculations

In order to describe the motion of the rigid bodies in our system, we have to obtain their corresponding diffusion tensors. Since both the NLSJ and the cargo are spheres (although we still treat them as rigid bodies with an internal OFR), their diffusion tensors are easily obtained. The 3 × 3 translational component of the diffusion tensor of a sphere is 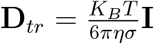, where *σ* is the sphere radius and **I** is the identity matrix. Similarly, the rotational component of the tensor is 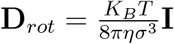. Since there is no coupling between the translation and rotation of a sphere, the coupling component of the tensor, **D**_*c*_, is a 3 × 3 null matrix.

Due to the complexity of the shape of the kinesin motor domain, it cannot be treated as a sphere and its diffusion tensor has to be obtained by more elaborate means. Given, a hydrodynamic interactions tensor **H**, the procedure for obtaining the 6 × 6 diffusion tensor has been described in detail by Bernal *et al* .^47^ For the hydrodynamic interactions, we use the Rotne-Prager-Yamakawa tensor,^48,49^

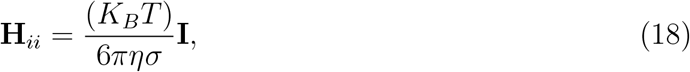

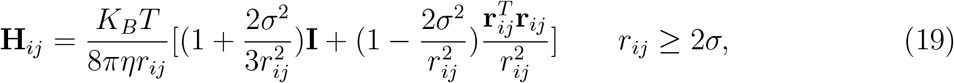

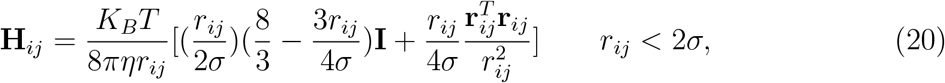

where **r**_*ij*_ is the vector connecting the *C*_*α*_ carbons of amino acids *i* and *j* in the motor domain, *r*_*ij*_ = |**r**_*ij*_|, and *σ* is the amino acid radius *σ*_*AA*_.

### Parametrization of Interactions

#### MT Binding

The kinesin step is dominated by two energy scales: the interactions energy, *ϵ*_*MT*_, between the motor domains and the MT, and the interaction energy, *ϵ*_*d*_, that governs docking.^26^ The value of the MT binding energy parameter *ϵ*_*MT*_ depends on the nucleotide state of the motor domain in question. The *apo* and ATP states have high affinity for MT, whereas the ADP state has low affinity for MT. ^35^ To obtain an estimate of the appropriate value of *ϵ*_*MT*_ in each of the nucleotide states, we performed unbinding simulations of a single motor domain bound to MT where an increasing external force was applied to the cargo.

The mean unbinding force of a single headed kinesin in different nucleotide states has been determined by Uemura *et al* .^35^ To determine the value of *ϵ*_*MT*_ in the high affinity state, we used the mean unbinding force data of kinesin bound to AMP-PNP, with the resistive force applied in the direction of the MT minus end. Similarly, the value of *ϵ*_*MT*_ in the low affinity state was determined using the data of the kinesin bound to ADP, with the assiting force applied in the direction of the MT plus end. The choice of force directions was made to account for the fact that the LH is predominantly in the high MT affinity state, whereas the TH is the motor domain that transitions to the low MT affinity state.

The experimental unbinding force studies were performed with a loading rate of *v* ≈ 5*pN/s*.^35^ This loading rate is too slow for our simulations, which are limited to millisecond timescales. In order to observe unbinding within a reasonable simulation time, we used a loading rate of *v* = 50*pN/µs*, which is seven orders of magnitude larger than the experimental value. Because the mean unbinding force depends on the loading rate logarithmically, *f*_*u*_ ∝ log (*v*),^50^ the estimates from the simulations could yield physical values (see the results in Fig. 6). We estimate the corresponding high and low affinity unbinding forces of the loading rate used in the simulations to be 43.5*pN* and 27.6*pN* respectively.

**Figure 6:**
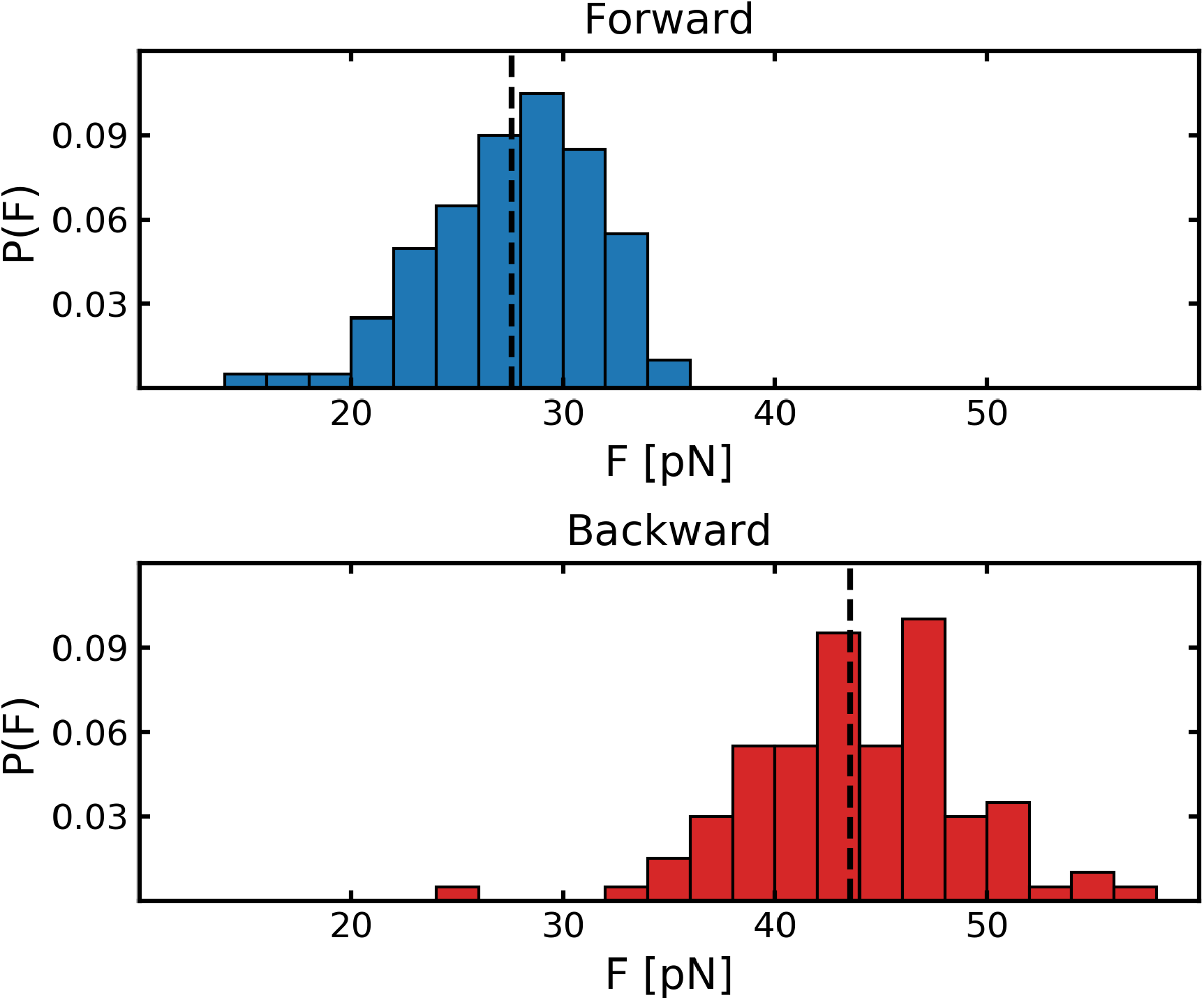
Top: A histogram of the unbinding forces of the TH when pulled forward toward the plus end of the MT. The expected mean unbinding force that was derived from experiment is indicated by the black dashed line. Bottom: The distribution of unbinding forces of the LH when pulled backward toward the minus end of the MT. The dashed line indicates the expected mean unbinding force, obtained from experiment.

As we mentioned earlier, we determined the values of the MT binding energy parameter to be 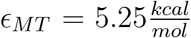 in the high affinity state and 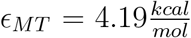 in the low affinity state. These values yield mean unbinding forces of 43.9 ± 0.5*pN* and 27.6 ± 0.4*pN*, respectively, which are similar to the experimentally derived forces. The distributions of unbinding forces, as well as the target mean unbinding forces are shown in Fig.6. As can be seen from the figure and from the calculated mean values, the simulations agree well with experiment, thus validating our choice of the *ϵ*_*MT*_ values.

#### NL Docking

To the best of our knowledge, there is no direct measurement of the energetics or kinetics of the NL docking process. It is, therefore, impossible to determine the appropriate value of *ϵ*_*d*_ with certainty. It is, however, possible to estimate a lower bound using the following argument. The mean run length ⟨*L*⟩ of kinesin is approximately ⟨*L*⟩ ≈ 1*µm*. The step size has been determined to be Δ*x* ≈ 8.1*nm*, corresponding to the distance between two adjacent *α/β* tubulin dimers in the MT.^6,51–53^ It is therefore possible to obtain the mean number of steps per run, 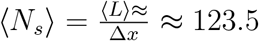. The mean number of steps has a direct dependence on the probability of detachment per step. If *P*_*d*_ is the probability of detachment, and *n* is the number of steps, than the mean number of steps per run is given by,

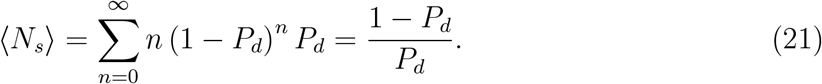

In other words, the probability that the motor steps *n* times is the probability that the motor does not detach during the first *n* steps, and detaches during step *n* + 1. Since we know the value of ⟨*N*_*s*_⟩ = ⟨*L* ⟩/Δ*x*, we can use Eq.21 to obtain the probability of detachment per step, 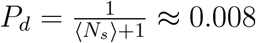.

We assume that the termination of the walk along the MT is primarily due to detachment of the motor from the MT during the step itself. This is a reasonable assumption because during the waiting time between steps, both motor domains are tightly bound to MT. Therefore, in all likelihood the walk is terminated when the motor is in the vulnerable one head bound state. During the step, there are two competing processes: detachment of the bound LH, and completion of the step. The detachment process is governed by the kinetics of ATP hydrolysis. Both in vivo, and in the experimental setups in which the run lengths are measured, the concentration of ATP is high and close to saturation. We assume that the LH is either bound to ATP, or binds it immediately at the beginning of the step. Therefore, the detachment rate of the bound LH is determined by the amount of time it takes the LH to hydrolyze the ATP molecule, and release the inorganic phosphate. The typical time for this process, which corresponds to the dwell time at high ATP concentrations, is approximately *τ*_*d*_ = 0.01*s*.^10^

Completion of the step requires two events. The first is for the diffusing TH to reach the TBS. The second is for the TH to release the ADP molecule once it reaches (or prior to reaching) the TBS, which in turn stabilizes the interaction with the MT. The kinetics of the former process is governed by the kinetics of the NL docking as well as the diffusive properties of the motor domain. The latter is determined by the release rate of the ADP when the motor is bound to the MT. We do not know the rate of ADP release, though it has been shown that it is greatly accelerated when the motor is bound to MT.^54^ We assume that this rate is significantly faster than the rate at which the TH reaches the TBS. Under this assumption, the time to complete the step is governed predominantly by the docking time.

To determine the stepping time, we simulated multiple trajectories of the kinesin step and measured the probability that the TH reaches the TBS as a function of time (see Fig.7). To fit our data, we used the cumulative probability function for a series of *n* consecutive Poisson processes,

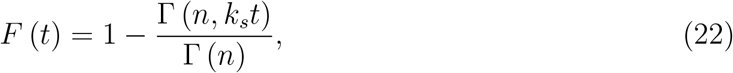

with *n* = 3, and *k*_*s*_ = 0.037*µs*^*−*1^, where Γ (*n, k*_*s*_*t*) is the incomplete gamma function. The goal in choosing this functional form and parameter values was purely to obtain an analytic function that fits our results well. As such, it should not be assigned an physical interpretation. The corresponding probability function is given by,

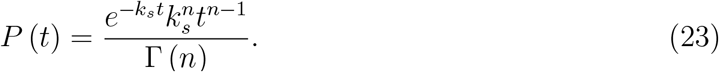

**Figure 7:**
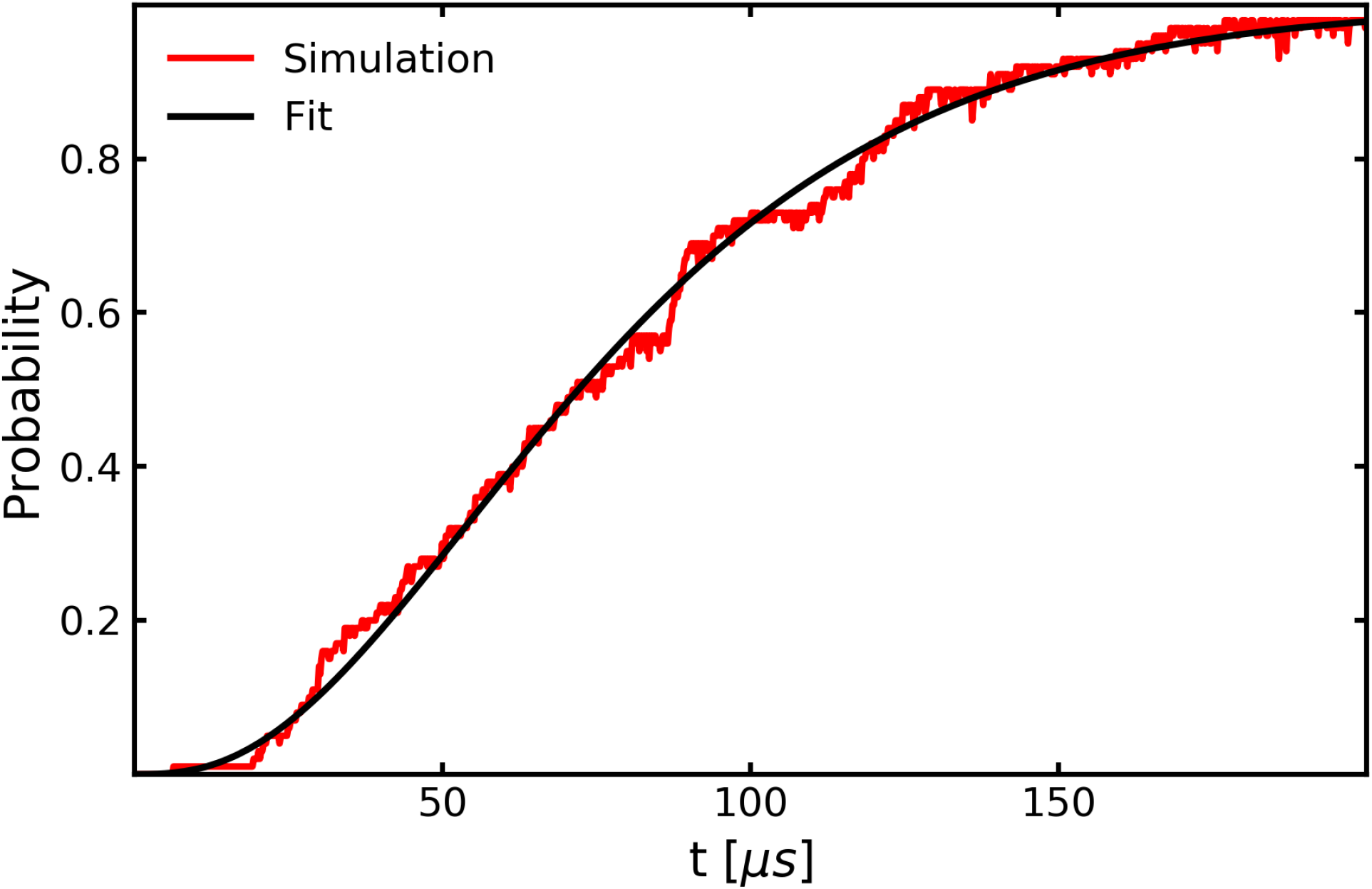
The probability that the TH binds to the TBS as a function of time (red). The data was fit using the cumulative probability density function shown in Eq.22.

If 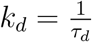 is the detachment rate of the LH, the detachment probability is given by,

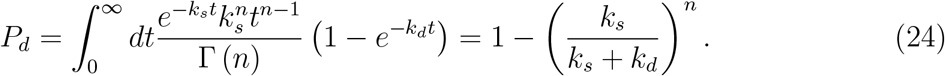

When *n* = 3, and *k*_*s*_ = 0.037*µs*^*−*1^, we obtain *P*_*d*_ ≈ 0.008 which is the value we expect according to the experimental findings. These values were obtained when we used 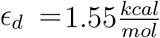 as the value for the docking energy parameter.

It is important to note that this result is valid only if the ADP release rate is much greater than the rate of completing the step. If that is not the case, the step completion kinetics will also depend on the ADP release rate. In such a scenario, faster docking is allowed, and as a consequence, higher values of *ϵ*_*d*_ are needed, as docking is no longer the rate limiting step in the process. Therefore, 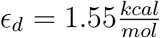 is the lower bound on the docking energy parameter.

## Results

### Docking Confines the TH Diffusive Search Space

With the calibrated model, we performed multiple simulations to obtain quantitative insights into the dynamics of a single step. In particular, we can quantify the contribution of the NL docking to the stepping process. We generated two sets of 50 trajectories of the complete kinesin step using two values of *ϵ*_*d*_ (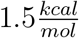 and 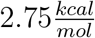), and monitored the position of the TH throughout the simulations. We projected the position of the TH onto the xy plane to give us a bird’s eye view of the trajectory. The x axis, referred to as the on axis, lies along the MT’s main axis. The y axis is the off axis along the plane on which the MT lies. The results are shown in Fig.8.

**Figure 8:**
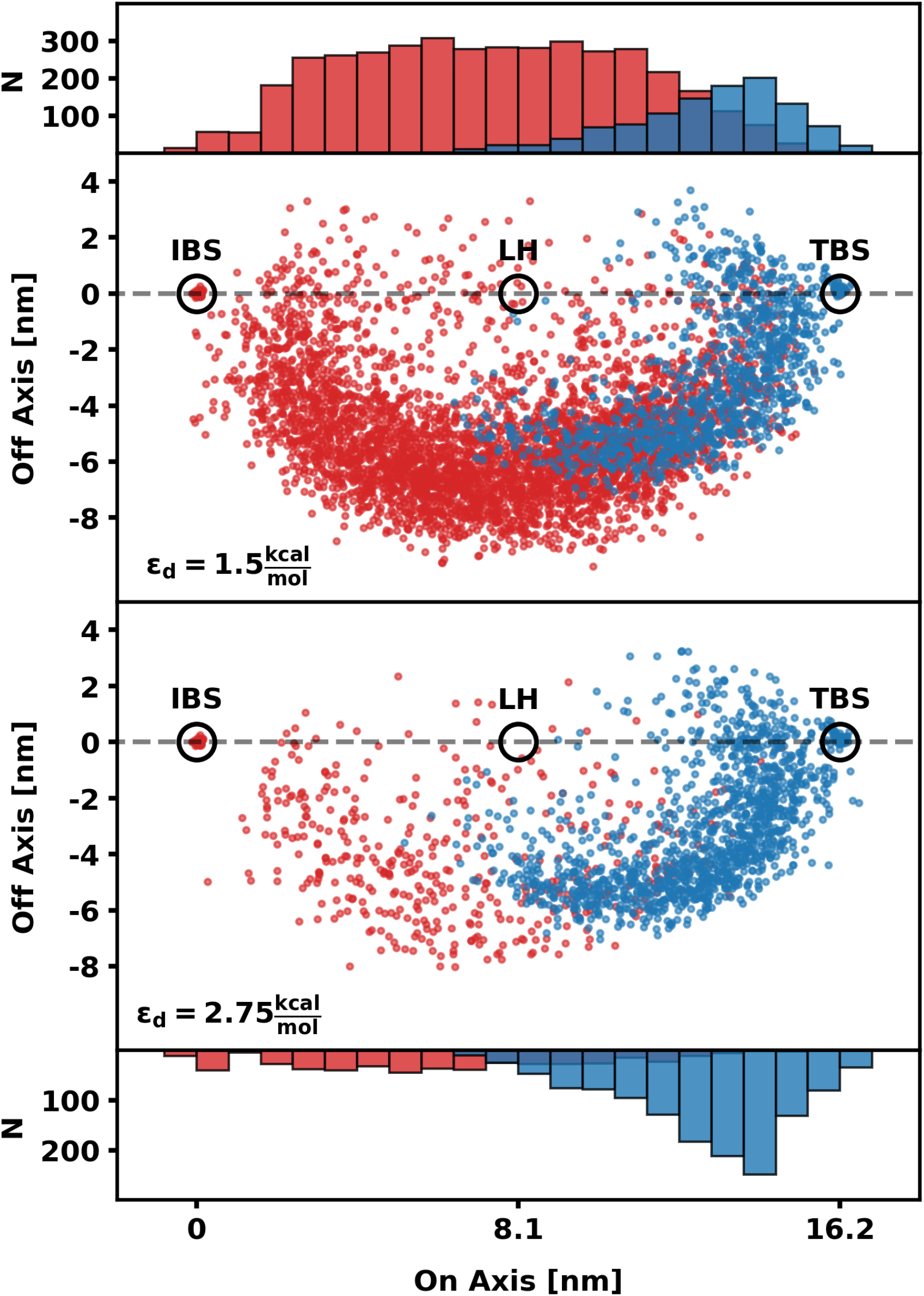
Positions of the TH projected onto the xy plane during the stepping process. The red (blue) points show the positions of the TH when the NL is undocked (docked). The Initial Binding Site (IBS) of the TH before detachment, as well as the positions of the LH and the TBS are indicated by black circles. The histograms correspond to the distribution along the x axis of the position of the TH during the step. Similarly to the middle panels, red and blue histograms correspond to the undocked and docked data points respectively. The two upper panels corresponds to simulations with 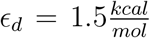 while the two lower panels corresponds to simulations with 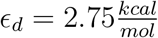.

To asses the contribution of the NL docking to the step, we highlighted the position of the TH when the NL is docked (blue data points in Fig.8). Our simulations show that when the NL is docked to the LH, the motion of the TH is confined to the space near the TBS (upper panels in Fig.8). When the NL is not docked, the TH explores a larger region around the LH, predominantly on its right side. This is supported by the observations made by Isojima *et al*. where the position of the TH was tracked using a gold nano-particle.^10^ Additionally, this result is consistent with the recent findings by Sudhakar *et al*., where at low ATP concentration (a condition under which we would expect a low probability of docking) the motor is shown to take sub steps, implying a possible temporary side step of the motor.^55^ Although the TH can reach the neighborhood of the TBS even without NL docking to the LH, it does so with a smaller probability when compared with the situation in which the NL is in the docked state. A comparison between the two sets of simulations (top and bottom panels in Fig.8) shows that increasing the value of *ϵ*_*d*_ shifts the balance towards the docked state, thus significantly increasing the probability that the TH will reach the TBS.

Given a long enough time, the TH would reach the TBS regardless of the NL docking. The NL docking is not technically required to complete the step, however, it does reduce the amount of time before the step is complete. To quantify the effect of the docking process on the stepping kinetics, we performed multiple sets of simulations with different values of *ϵ*_*d*_ and measured the first passage time for docking of the NL as well as the time required to complete the step (see SI for details).

Our results (Fig.9) show that as *ϵ*_*d*_ increases, the mean docking and stepping time decrease accordingly. It is worth noting that the docking and stepping times plateau at the higher values of *ϵ*_*d*_, with the completion of the step taking place approximately 5*µs* after docking is complete. The reason for this is that the stepping process is governed and limited by the diffusive motion of the TH.^34^ Therefore, even if the docking process was instantaneous, the TH would take some time to diffuse before reaching the TBS. This is vividly illustrated in Fig.8. While the docked NL limits the search space of the TH, it still has to diffuse within the confined region before reaching the TBS. The lowest value of *ϵ*_*d*_ that we used was 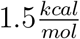, just below the lower bound of 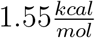. At this value the step was complete within 20*µs*. We can therefore estimate that the stepping process is completed within the range of 5-20*µs*, which in accord with experimental estimates.

**Figure 9:**
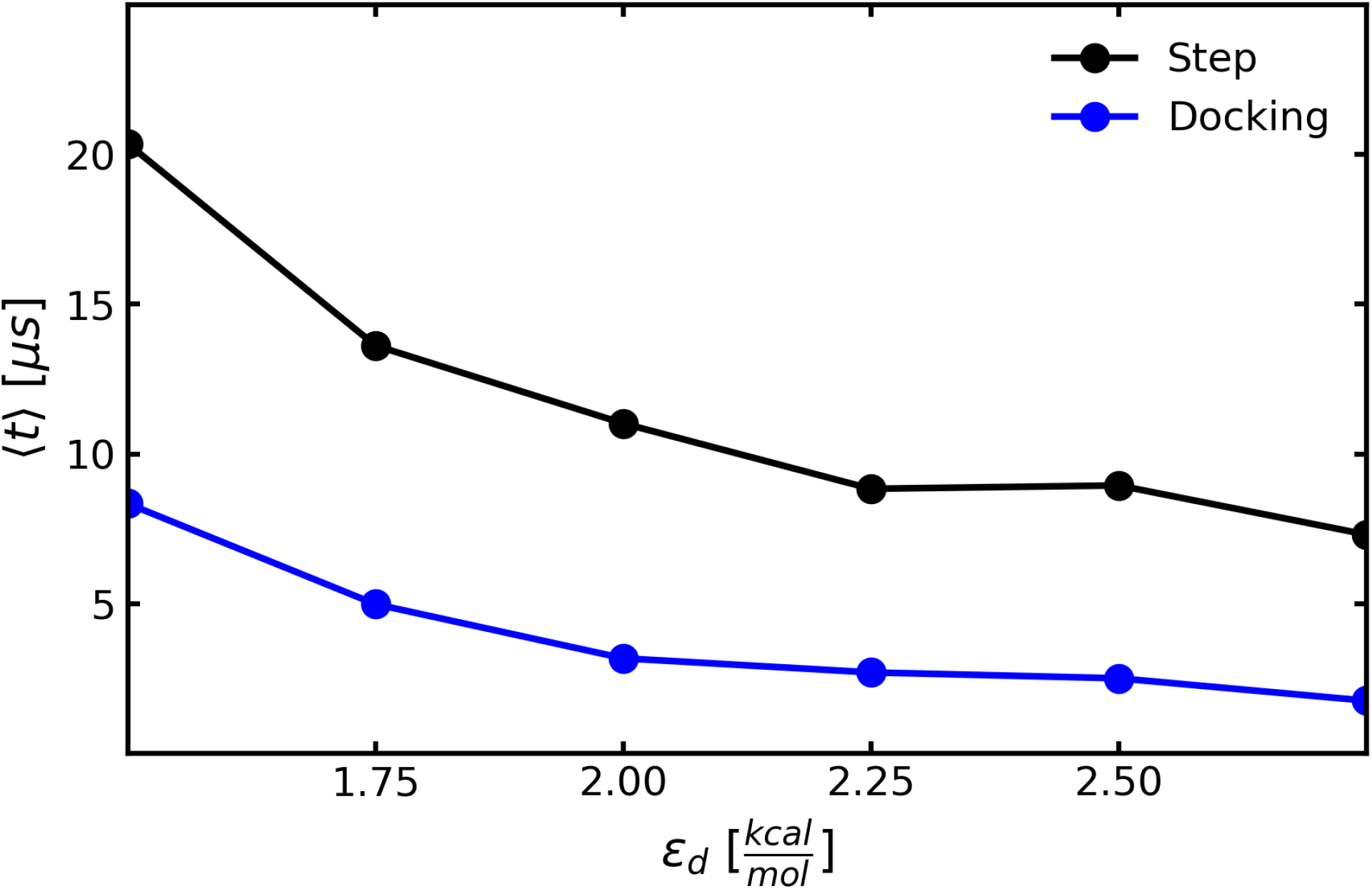
First passage time for the docking of the NL (blue), as well as the time to complete the step (black) as a function of the value of the docking energy parameter *ϵ*_*d*_.

### Torsional Tension along the Stalk

A free parameter in our model is the torsional spring constant *k* in Eq.7, which we will refer to as *k*_*stalk*_. This parameter determines the torsional flexibility of the stalk which plays an important role in the interaction of the motor with the cargo. The value of this parameter depends on the nature of the connection of the stalk with the cargo as well as the tensile properties of the stalk itself. It has been shown that different stalk constructs result in different levels of torsional stiffness.^40^ The torsional potential in Eq.7 plays an important role in the torsional constraint that the motor imposes on the cargo (and vice versa). Furthermore, the torsional stiffness is also influenced by the torque that is exerted by the two NLs at the NLSJ.

In order to determine the effective stiffness of the torsional constraint, it is necessary to simulate the two heads bound state and track the rotational movement of the cargo. Due to its large size, the rotational diffusion of the cargo is a slow process, occurring on a time scale that is significantly longer than our simulation time scale. To circumvent this problem, we artificially decreased the z component of the cargo friction tensor to accelerate the cargo rotational dynamics along the z axis of the OFR. Since we are interested in an equilibrium property, namely the effective torsional spring constant, our result is independent of the value of the friction tensor.

We performed multiple simulation sets, consisting of 100 trajectories each, using different values of *k*_*stalk*_. We then measured the angular motion of the cargo, obtained the angular distribution, and fitted it with a Gaussian distribution to extract the corresponding effective spring constant *k*_*eff*_. The value of *k*_*eff*_ as a function of *k*_*stalk*_ is shown in Fig.10.

**Figure 10:**
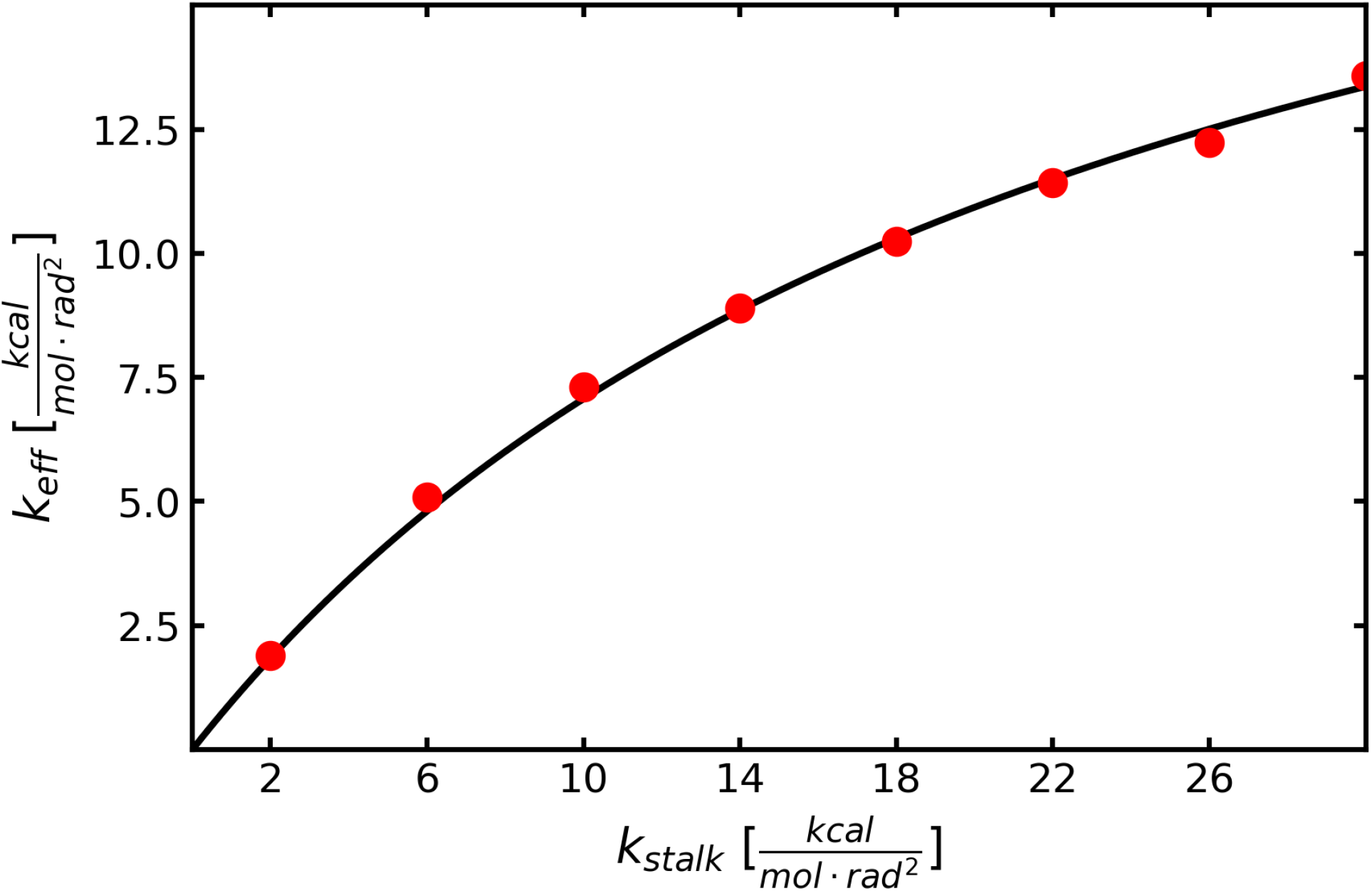
The effective torsional stiffness spring constant *k*_*eff*_ as a function of the stalk torsional spring constant *k*_*stalk*_ (red). The data points were fitted using the expression for the effective spring constant in Eq.26 (black).

As expected, *k*_*eff*_ increases as the value of *k*_*stalk*_ increases, however, at higher values of *k*_*stalk*_ it appears to plateau. It is reasonable to suspect that *k*_*eff*_ has an upper bound because even if the stalk was infinitely stiff, the NLs would allow for some torsional flexibility. In order to estimate the upper bound of *k*_*eff*_ we used the following simplification. The total torsional energy can be expressed as,

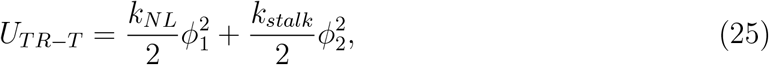

where *ϕ*_1_ represents the rotation of the NLSJ relative to the MT axis, and *ϕ*_2_ represents the internal torsional rotation of the stalk. The spring constant *k*_*NL*_ accounts for the rotational tension acting on *ϕ*_1_ due to the tension along the NLs. The second term in Eq.25 corresponds to the torsional energy in Eq.7, whereas the first term rises from the connectivity and excluded volume terms in the Hamiltonian.

The overall cargo rotation can be expressed in terms of Φ = *ϕ*_1_ + *ϕ*_2_. We can express Eq.25 in terms of ϕ. We assume that the dynamics of the NLs takes place on a time scale that is significantly shorter than that of the cargo rotation. This assumption is valid as long as we do not greatly accelerate the rotational dynamics of the cargo. In that case, we can assert that *ϕ*_1_ and *ϕ*_2_ reach a mechanical equilibrium before any appreciable change in ϕ takes place. If so, on average, *ϕ*_1_ assumes a value that minimizes the energy in Eq.25. This results in the following approximation of the torsional energy,

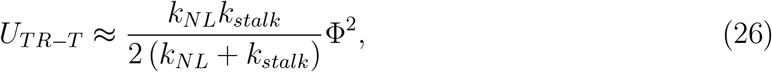

where the effective spring constant is 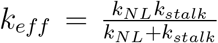. We fit the expression for *k*_*eff*_ to our data points from the simulations (black line in Fig.10), and determined the value of 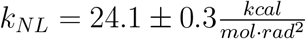, which is also the upper bound of *k*_*eff*_ .

To further investigate the tensile properties of kinesin, we calculated the tension exerted on the TH by the LN in the two head bound state. We first calculated the magnitude of the mean force that is exerted on the TH rigid body by the FENE potential at the point of contact with the NL and obtained a force of *F* ≈ 60*pN*. Our force estimate is larger than previous estimates which place the force at around *F* ≈ 12 − 15*pN* .^56^ In all likelihood the force obtained in our simulations is an overestimate of the actual force. We suspect that this is due to the rigid nature of the motor domain as well as the NLSJ, which does not allow for any flexibility at the contact points with the NLs. Since the NLs may be modeled as worm-like chains, any over-extension, even if small, would lead to a larger force. It is important to note that, for this reason, our predicted value of the upper bound for the torsional stiffness may be an overestimate as well. To determine whether this is actually the case, the upper bound for the torsional stiffness can be tested experimentally.

## Conclusions

We have created a new multi-scale coarse-grained model, consisting of rigid and flexible elements of different sizes, that can be used to efficiently simulate the stepping mechanism of the kinesin motor. The main advantage of our approach is the computational efficiency, which allows us to generate multiple trajectories for a single step, and allows us to answer questions that cannot be addressed by any other computational model. Interestingly, despite the high level coarse graining used to derive the MCSG, our model captures many important aspects of the kinesin step without any significant loss in accuracy. Due to high computational efficiency, it is possible to fit experimental data systematically to ensure the validity and accuracy of the model. Because the model has only few parameters, one could explore the parameter space in order to investigate how the behavior of the motor would change if the parameters are varied, which is useful in the design of artificial motors.

The rigid body model of kinesin opens the door to explore the behavior of the motor over multiple steps. In particular, the effects of the cargo on the stepping mechanism is an important open question that is yet to be fully resolved in experiments. Furthermore, this approach facilitates the simulation of multiple motors at the same time, which is relevant to biology, since cargo transport in vivo if often carried out by many (different) motors that are bound to the same cargo.

A second important advantage of our approach is the modularity, which allows for a flexible design of the model that is tailored to other motors. The main limitation is that the rigid body formulation can only be used for elements that do not undergo large conformational changes during the process of interest. Any flexible or deformable component has to be modeled accordingly, as we have done for the NLs and the stalk in the case of kinesin. We believe that the MSCG model would be particularly useful in the description of the behavior of dynein and myosin.

## Acknowledgments

This work was supported in part by a grant from National Science Foundation (CHE 19-00093) and the Collie-Welch Chair (F-0019) administered through the Welch Foundation.

## References

(1) Hirokawa, N.; Noda, Y.; Tanaka, Y.; Niwa, S. Kinesin superfamily motor proteins and intracellular transport. Nature reviews Molecular cell biology 2009, 10, 682–696.

(2) Hammer, J. A.; Sellers, J. R. Walking to work: roles for class V myosins as cargo transporters. Nature Reviews Molecular Cell Biology 2012, 13, 13–26.

(3) Roberts, A. J.; Kon, T.; Knight, P. J.; Sutoh, K.; Burgess, S. A. Functions and mechanics of dynein motor proteins. Nature reviews Molecular cell biology 2013, 14, 713–726.

(4) Mugnai, M. L.; Hyeon, C.; Hinczewski, M.; Thirumalai, D. Theoretical perspectives on biological machines. Reviews of Modern Physics 2020, 92, 025001.

(5) Yildiz, A.; Forkey, J. N.; McKinney, S. A.; Ha, T.; Goldman, Y. E.; Selvin, P. R. Myosin V walks hand-over-hand: single fluorophore imaging with 1.5-nm localization. science 2003, 300, 2061–2065.

(6) Yildiz, A.; Tomishige, M.; Vale, R. D.; Selvin, P. R. Kinesin walks hand-over-hand. Science 2004, 303, 676–678.

(7) Reck-Peterson, S. L.; Yildiz, A.; Carter, A. P.; Gennerich, A.; Zhang, N.; Vale, R. D. Single-molecule analysis of dynein processivity and stepping behavior. Cell 2006, 126, 335–348.

(8) DeWitt, M. A.; Chang, A. Y.; Combs, P. A.; Yildiz, A. Cytoplasmic dynein moves through uncoordinated stepping of the AAA+ ring domains. Science 2012, 335, 221–225.

(9) Qiu, W.; Derr, N. D.; Goodman, B. S.; Villa, E.; Wu, D.; Shih, W.; Reck-Peterson, S. L. Dynein achieves processive motion using both stochastic and coordinated stepping. Nature structural & molecular biology 2012, 19, 193–200.

(10) Isojima, H.; Iino, R.; Niitani, Y.; Noji, H.; Tomishige, M. Direct observation of intermediate states during the stepping motion of kinesin-1. Nature chemical biology 2016,

(11) Kull, F. J.; Sablin, E. P.; Lau, R.; Fletterick, R. J.; Vale, R. D. Crystal structure of the kinesin motor domain reveals a structural similarity to myosin. Nature 1996, 380, 550.

(12) Sack, S.; Müller, J.; Marx, A.; Thormählen, M.; Mandelkow, E.-M.; Brady, S. T.; Mandelkow, E. X-ray structure of motor and neck domains from rat brain kinesin. Biochemistry 1997, 36, 16155–16165.

(13) Shang, Z.; Zhou, K.; Xu, C.; Csencsits, R.; Cochran, J. C.; Sindelar, C. V. Highresolution structures of kinesin on microtubules provide a basis for nucleotide-gated force-generation. Elife 2014, 3, e04686.

(14) Carter, A. P.; Cho, C.; Jin, L.; Vale, R. D. Crystal structure of the dynein motor domain. Science 2011, 331, 1159–1165.

(15) Kon, T.; Oyama, T.; Shimo-Kon, R.; Imamula, K.; Shima, T.; Sutoh, K.; Kurisu, G. The 2.8 Å crystal structure of the dynein motor domain. Nature 2012, 484, 345–350.

(16) Schmidt, H.; Gleave, E. S.; Carter, A. P. Insights into dynein motor domain function from a 3.3-Å crystal structure. Nature structural & molecular biology 2012, 19, 492–497.

(17) Whitford, P. C.; Sanbonmatsu, K. Y.; Onuchic, J. N. Biomolecular dynamics: order-disorder transitions and energy landscapes. Rep. Prog. Phys. 2012, 75, 076601.

(18) Hyeon, C.; Thirumalai, D. Capturing the essence of folding and functions of biomolecules using coarse-grained models. Nature communications 2011, 2, 487.

(19) Hyeon, C.; Onuchic, J. N. Mechanical control of the directional stepping dynamics of the kinesin motor. Proc. Natl. Acad. Sci. U. S. A. 2007, 104, 17382–17387.

(20) Hyeon, C.; Lorimer, G. H.; Thirumalai, D. Dynamics of allosteric transition in GroEL. Proc. Natl. Acad. Sci. U. S. A. 2006, 103, 18939–18944.

(21) Koga, N.; Takada, S. Folding-based molecular simulations reveal mechanisms of the rotary motor F_1_-ATPase. Proc. Natl. Acad. Sci. U. S. A. 2006, 103, 5367–5372.

(22) Hori, N.; Takada, S. Coarse-Grained Structure-Based Model for RNA-Protein Complexes Developed by Fluctuation Matching. J. Chem. Theory Comp. 2012, 8, 3384–3394.

(23) Tanaka, T.; Hori, N.; Takada, S. How Co-translational Folding of Multidomain Protein Is Affected by Elongation Schedule: Molecular Simulations. PLOS Comp. Biol. 2015, 11, e1004356.

(24) Hyeon, C.; Onuchic, J. N. Mechanical control of the directional stepping dynamics of the kinesin motor. Proceedings of the National Academy of Sciences 2007, 104, 17382–17387.

(25) Tehver, R.; Thirumalai, D. Rigor to post-rigor transition in myosin V: link between the dynamics and the supporting architecture. Structure 2010, 18, 471–481.

(26) Zhang, Z.; Thirumalai, D. Dissecting the kinematics of the kinesin step. Structure 2012, 20, 628–640.

(27) Goldtzvik, Y.; L, M. M.; Thirumalai, D. Dynamics of Allosteric Transitions in Dynein. Structure 2018, 26, 1664–1677.

(28) Schnitzer, M. J.; Block, S. M. Kinesin hydrolyses one ATP per 8-nm step. Nature 1997, 388, 386.

(29) Hua, W.; Young, E. C.; Fleming, M. L.; Gelles, J. Coupling of kinesin steps to ATP hydrolysis. Nature 1997, 388, 390.

(30) Bhabha, G.; Cheng, H.-C.; Zhang, N.; Moeller, A.; Liao, M.; Speir, J. A.; Cheng, Y.; Vale, R. D. Allosteric communication in the dynein motor domain. Cell 2014, 159, 857–868.

(31) Mugnai, M. L.; Thirumalai, D. Kinematics of the lever arm swing in myosin VI. Proceedings of the National Academy of Sciences 2017, 114, E4389–E4398.

(32) Rice, S.; Lin, A. W.; Safer, D.; Hart, C. L.; Naber, N.; Carragher, B. O.; Cain, S. M.; Pechatnikova, E.; Wilson-Kubalek, E. M.; Whittaker, M. et al.. A structural change in the kinesin motor protein that drives motility. Nature 1999, 402, 778–784.

(33) Goldtzvik, Y.; Zhang, Z.; Thirumalai, D. Importance of Hydrodynamic Interactions in the Stepping Kinetics of Kinesin. The Journal of Physical Chemistry B 2016, 120, 2071–2075.

(34) Zhang, Z.; Goldtzvik, Y.; Thirumalai, D. Parsing the roles of neck-linker docking and tethered head diffusion in the stepping dynamics of kinesin. Proceedings of the National Academy of Sciences 2017, 114, E9838–E9845.

(35) Uemura, S.; Kawaguchi, K.; Yajima, J.; Edamatsu, M.; Toyoshima, Y. Y.; Ishiwata, S. Kinesin–microtubule binding depends on both nucleotide state and loading direction. Proceedings of the National Academy of Sciences 2002, 99, 5977–5981.

(36) Carter, N. J.; Cross, R. Mechanics of the kinesin step. Nature 2005, 435, 308–312.

(37) Guydosh, N. R.; Block, S. M. Backsteps induced by nucleotide analogs suggest the front head of kinesin is gated by strain. Proceedings of the National Academy of Sciences 2006, 103, 8054–8059.

(38) Valdenebro, A. G. Visualizing rotations and composition of rotations with the Rodrigues vector. Euro. J. Phys. 2016, 37, 065001.

(39) Wolgemuth, C. W.; Sun, S. X. Elasticity of α-helical coiled coils. Physical review letters 2006, 97, 248101.

(40) Gutiérrez-Medina, B.; Fehr, A. N.; Block, S. M. Direct measurements of kinesin torsional properties reveal flexible domains and occasional stalk reversals during stepping. Proceedings of the National Academy of Sciences 2009, 106, 17007–17012.

(41) Ramaiya, A.; Roy, B.; Bugiel, M.; Schäffer, E. Kinesin rotates unidirectionally and generates torque while walking on microtubules. Proceedings of the National Academy of Sciences 2017, 114, 10894–10899.

(42) Ermak, D. L.; McCammon, J. Brownian dynamics with hydrodynamic interactions. The Journal of chemical physics 1978, 69, 1352–1360.

(43) Leimkuhler, B.; Matthews, C. Rational construction of stochastic numerical methods for molecular sampling. Applied Mathematics Research eXpress 2012, 2013, 34–56.

(44) Antosiewicz, J.; Grycuk, T.; Porschke, D. Brownian dynamics simulation of electrooptical transients for solutions of rigid macromolecules. The Journal of chemical physics 1991, 95, 1354–1360.

(45) Gibbs, J. Vector analysis. A text-book for the use of students of mathematics and physics; Yale University Press, 2008.

(46) Evans, D. J.; Murad, S. Singularity free algorithm for molecular dynamics simulation of rigid polyatomics. Molecular physics 1977, 34, 327–331.

(47) Bernal, J. M. G., De La Torre, J. G. Transport properties and hydrodynamic centers of rigid macromolecules with arbitrary shapes. Biopolymers: Original Research on Biomolecules 1980, 19, 751–766.

(48) Rotne, J.; Prager, S. Variational treatment of hydrodynamic interaction in polymers. The Journal of Chemical Physics 1969, 50, 4831–4837.

(49) Yamakawa, H. Transport properties of polymer chains in dilute solution: hydrodynamic interaction. The Journal of Chemical Physics 1970, 53, 436–443.

(50) Evans, E.; Ritchie, K. Dynamic Strength of Molecular Adhesion Bonds. Biophysical journal 1997, 72, 1541–1555.

(51) Yildiz, A.; Tomishige, M.; Gennerich, A.; Vale, R. D. Intramolecular strain coordinates kinesin stepping behavior along microtubules. Cell 2008, 134, 1030–1041.

(52) Milic, B.; Andreasson, J. O.; Hancock, W. O.; Block, S. M. Kinesin processivity is gated by phosphate release. Proceedings of the National Academy of Sciences 2014, 111, 14136–14140.

(53) Andreasson, J. O.; Milic, B.; Chen, G.-Y.; Guydosh, N. R.; Hancock, W. O.; Block, S. M. Examining kinesin processivity within a general gating framework. Elife 2015, 4, e07403.

(54) Hackney, D. D. Kinesin ATPase: rate-limiting ADP release. Proceedings of the National Academy of Sciences 1988, 85, 6314–6318.

(55) Sudhakar, S.; Abdosamadi, M. K.; Jachowski, T. J.; Bugiel, M.; Jannasch, A.; Schäffer, E. Germanium nanospheres for ultraresolution picotensiometry of kinesin motors. Science 2021, 371.

(56) Hyeon, C.; Onuchic, J. N. Internal strain regulates the nucleotide binding site of the kinesin leading head. Proceedings of the National Academy of Sciences 2007, 104, 2175–2180.

