## Supplementary Information for "Multi-Scale Coarse Grained Model for the Stepping of Molecular Motors with application to Kinesin"

Yonathan Goldtzvik* and D. Thirumalai*

*Department of Chemistry, University of Texas at Austin, Austin, TX 78705*

### Coarse-Grained Model Building

Our goal was to design a simplified representation of the kinesin motor domain that preserves the overall geometrical features while avoiding a detailed atomistic description, which would make simulations difficult. To this end, we represented the motor domain using a relatively small number of beads (39 beads) of varying sizes. The choices of bead positions and sizes were as follows. We first divided the amino acids in the motor domain into groups, based on which amino acids were in the same region of the motor. The group choices were done manually and are somewhat arbitrary. However, we expect that the particular amino acid group choice is not of great consequence for the results of our study, as long as the general shape of the motor domain is maintained. An example of one such amino acid group together with the resulting simplified coarse-grained representation are shown in Fig. S1. The radius of each bead in the coarse-grained representation is the radius of gyration of the positions of the $C_\alpha$ carbons of the amino acids in the group. A list of the amino acid groups and their respective radii can be found in Table S1. It is important to note that in order to maintain the flexibility of the Cover-Neck Bundle (CNB) and the Neck Linker (NL), we represented each of the amino acids in these structure as a single bead and did not include them in an amino acid group, unlike the rest of the motor domain.

### Interactions the Between Motor Domain and Microtubule

The interactions between the motor domain and the Microtubule (MT) binding sites in our model are described by the following energy term,

$$U_{MT} = -\sum_{i=1}^{6} \frac{\epsilon_{MT}}{1 + k\left(\mathbf{r}_i - \mathbf{r}_i^0\right)^2}, \tag{1}$$

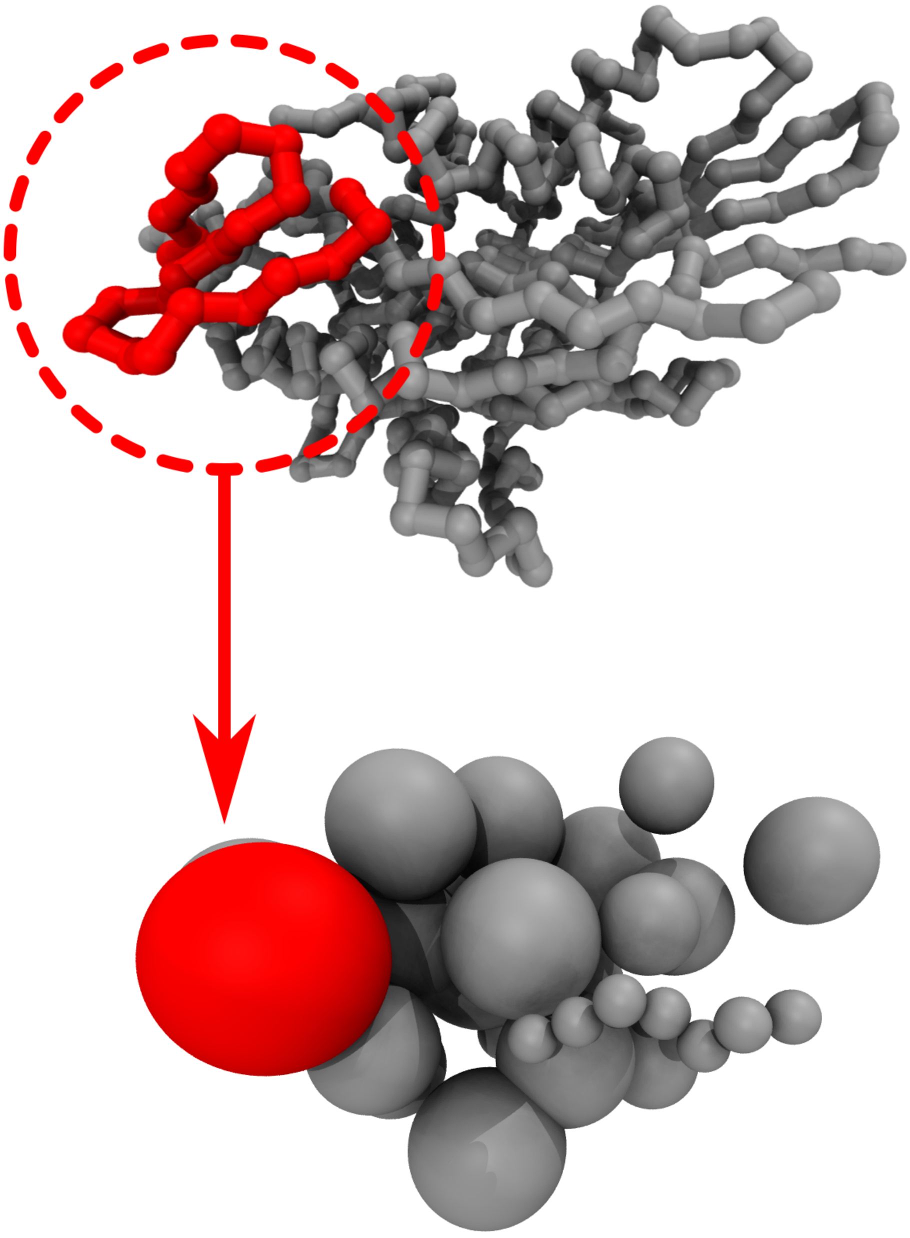

Figure S1: An example of one group of amino acids in the kinesin atomistic structure represented as a single coarse grained bead.

**Table S1: CG Model $C_\alpha$ Groups in Kinesin Structure**

| Group # | Residues | Radius |
|---|---|---|
| 1 | 1 | 1.9 |
| 2 | 2 | 1.9 |
| 3 | 3 | 1.9 |
| 4 | 4 | 1.9 |
| 5 | 5 | 1.9 |
| 6 | 6 | 1.9 |
| 7 | 7 | 1.9 |
| 8 | 15-28,291-300 | 7.66 |
| 9 | 29-50 | 7.64 |
| 10 | 51-63 | 6.28 |
| 11 | 7-10,64-70,80-82,285-287 | 6.92 |
| 12 | 71-79 | 4.49 |
| 13 | 11-14,83-94,288-290 | 6.52 |
| 14 | 95-114 | 6.55 |
| 15 | 115-122 | 4.25 |
| 16 | 123-127,215-222 | 5.66 |
| 17 | 128-130,210-214,223-227 | 5.81 |
| 18 | 131-134,170-184 | 6.87 |
| 19 | 135-136,143-154,165-169 | 6.49 |
| 20 | 137,142 | 3.55 |
| 21 | 155-164 | 5.0 |
| 22 | 185-209,228-233 | 9.6 |
| 23 | 234-246 | 5.63 |
| 24 | 247-269 | 7.37 |
| 25 | 270-284 | 6.73 |
| 26 | 301,312 | 5.65 |
| 27 | 313 | 1.9 |
| 28 | 314 | 1.9 |
| 29 | 315 | 1.9 |
| 30 | 316 | 1.9 |
| 31 | 317 | 1.9 |
| 32 | 318 | 1.9 |
| 33 | 319 | 1.9 |
| 34 | 320 | 1.9 |
| 35 | 321 | 1.9 |
| 36 | 322 | 1.9 |
| 37 | 323 | 1.9 |
| 38 | 324 | 1.9 |
| 39 | 325 | 1.9 |

where the summation is over the 6 beads in our coarse grained representation of the motor domain, and $\mathbf{r}_i^0$ in Eq. 1 corresponds to the position of bead $i$, based on the crystal structure of the bound state (PDBID: 2P4N). The beads that were involved in these interactions are those that are at the interface between the motor domain and the MT in the bound state (groups 18-20 and 22-24 in Table S1). Since the motor domain can bind to any $\alpha\beta$ tubulin dimer, each dimer has 6 pseudo bead positions to which the 6 beads in the motor domain can bind. This is illustrated in Fig. S2

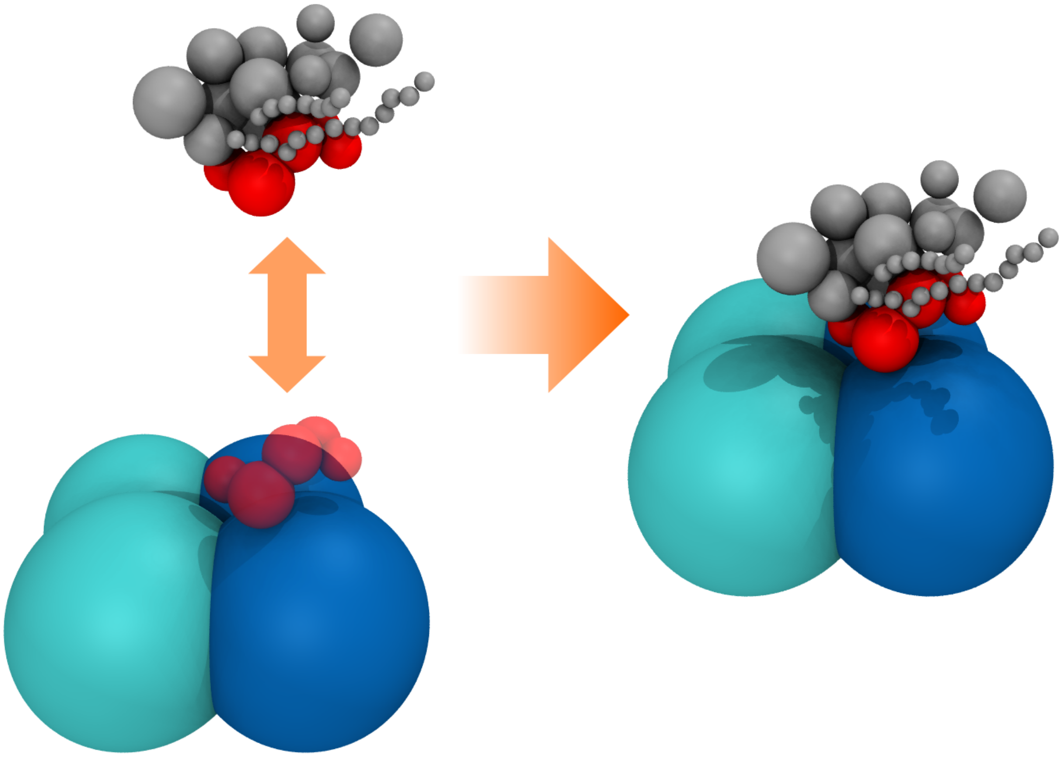

Figure S2: Graphical representation of the binding of the motor domain to the MT. The beads that correspond to the MT binding interface in the motor domain are highlighted in red. The correspondinging binding positions on the $\alpha\beta$ tubulin dimer are also highlighted.

### First Passage Times Calculations

In our analyses, we calculate the first passage times for the completion of the docking process, as well as the termination of the step (see main text). To do so, we first define a metric, which we is used to determine whether the NL of the Leading Head (LH) is docked and whether the Trailing Head (TH) reached the Target Binding Site (TBS). By defining a cutoff value for these metrics, we can determine whether the processes in question are complete.

We first address the docking of the NL of the LH. The NL has 13 amino acids of which only 12 are free to move in relation to the Object Frame of Reference (OFR; see Fig. 3 in the main text) of the LH. Each bead $i$, representing one of the 12 NL amino acids, has a low energy position within the OFR of the LH, $\mathbf{r}_i^0$, which corresponds to the position in the docked state (see Eq. 10 in the main text). We define the following metric,

$$\Delta_{NL} = \sqrt{\frac{\sum_i^N (\mathbf{r}_i - \mathbf{r}_i^0)^2}{N}}, \tag{2}$$

where the sum is over the $N = 12$ beads of the NL. This metric, $\Delta_{NL}$, measures the deviation of the NL from the docked state. We use a cutoff value of $\Delta_{NL} \leq 15\mathring{A}$ for the docked state.

In order to assess the binding of the TH to the TBS, we define the following metric. When bound to the TBS, the position of the TH is defined as $\mathbf{r}_{TH}^{TBS}$. Given that the position of the diffusing TH is given by $\mathbf{r}_{TH}$, the distance of the TH to the TBS is defined as,

$$\Delta_{TBS} = \sqrt{(\mathbf{r}_{TH} - \mathbf{r}_{TH}^{TBS})^2}. \tag{3}$$

We choose the value $\Delta_{TBS} < 3.8\mathring{A}$ as the cutoff for the bound state. To ensure that the TH is stably bound to the TBS, we require that $\Delta_{TBS} \leq 3.8\mathring{A}$ continuously for at least $0.2\mu s$.
